# Amyloid-beta peptides 40 and 42 employ distinct molecular pathways for cell entry and intracellular transit at the BBB endothelium

**DOI:** 10.1101/2022.11.17.516996

**Authors:** Zengtao Wang, Nidhi Sharda, Rajesh S. Omtri, Ling Li, Karunya K. Kandimalla

## Abstract

Blood-brain barrier (BBB) is a critical portal regulating the bidirectional transport of amyloid beta (Aβ) proteins between blood and brain. Disrupted trafficking at the BBB may not only promote the build-up of Aβ plaques in the brain parenchyma, but also facilitate Aβ accumulation within the BBB endothelium, which aggravates BBB dysfunction. Soluble Aβ42:Aβ40 ratios in plasma and cerebrospinal fluid have been reported to decrease during Alzheimer’s disease (AD) progression. Our previous publications demonstrated that trafficking of Aβ42 and Aβ40 at the BBB is distinct and is disrupted under various pathophysiological conditions. However, the intracellular mechanisms that allow BBB endothelium to differentially handle Aβ40 and Aβ42 have not been clearly elucidated. In this study, we identified mechanisms of fluorescently labeled Aβ (F-Aβ) endocytosis in polarized human cerebral microvascular endothelial (hCMEC/D3) cell monolayers using pharmacological inhibition and siRNA knock-down approaches. Further, intracellular transit of F-Aβ following endocytosis was tracked using live cell imaging. Our studies demonstrated that both F-Aβ peptides were internalized by BBB endothelial cells via energy, dynamin and actin dependent endocytosis. Interestingly, endocytosis of F-Aβ40 is found to be clathrin-mediated, whereas F-Aβ42 endocytosis is caveolae-mediated. Following endocytosis, both isoforms were sorted by the endo-lysosomal system. While Aβ42 was shown to accumulate more in the lysosome which could lead to its higher degradation and/or aggregation at lower lysosomal pH, Aβ40 demonstrated robust accumulation in recycling endosomes which may facilitate its transcytosis across the BBB. These results provide a mechanistic insight into the selective ability of BBB endothelium to transport Aβ40 versus Aβ42. This knowledge contributes to the understanding of molecular pathways underlying Aβ accumulation in the BBB endothelium and associated cerebrovascular dysfunction as well as amyloid deposition in the brain parenchyma which are implicated in AD pathogenesis.

## INTRODUCTION

Mounting evidence suggests that reduction in the ratio of amyloid beta peptide 42 (Aβ42) and Aβ40, two major Aβ isoforms that accumulate in AD patient plasma and the cerebrospinal fluid is associated with the severity of AD pathology and is emerging as a reliable biomarker for AD diagnosis^1–2^. Changes in Aβ42/Aβ40 ratio during AD progression may not be solely driven by the differential amyloidogenicity of Aβ42 and Aβ40; differential handling of Aβ0 and Aβ42 by the blood-brain barrier (BBB) and other disposition (distribution, metabolism, and elimination) pathways may also play a substantial role.

BBB is a physiological interface between blood and the brain which maintains the brain homeostasis by delivering essential nutrients to the brain and removing toxic metabolites out of the brain^3–4^. The carrier and barrier functions of the BBB are coordinated by selective internalization and intricate intracellular trafficking apparatus ^5^. The luminal-to-abluminal transport of Aβ was reported to occur via the receptor for advanced glycation end products (RAGE)^6^, whereas the low-density lipoprotein receptor related protein-1 (LRP1)^7–8^ and P-glycoprotein (P-gp)^9^ were claimed to mediate brain-to-blood efflux. However, the molecular machinery by which these receptors/transporters mediate cell entry and subsequent intracellular transit of Aβ in the BBB endothelial cells is only partially characterized.

Impairment of Aβ transport at the BBB, is believe to promote the formation of cerebrovascular amyloid deposits and amyloid plaques found in the brains of Alzheimer’s disease (AD) patients. Disrupted Aβ trafficking at the BBB is also presumed to increase anomalous accumulation of Aβ peptides in the BBB endothelium that compromise BBB integrity^10^ and function^11^, and trigger inflammatory changes^12^. Therefore, elucidating molecular mechanisms underlying Aβ trafficking disruption is critical for identifying novel therapeutic strategies.

Our recent study demonstrated that these two isoforms exhibit distinct trafficking and accumulation kinetics at the BBB endothelium^13^, which could be attributed to distinct trafficking apparatus employed by Aβ40 and Aβ42, although they are thought to be transported by the same receptors/transporters. Elucidation of the underlying mechanisms by which Aβ isoforms are differentially internalized and sorted in the BBB endothelial cells may lead to the identification of molecular mediators that drive pathological shifts in Aβ42/40 ratios in plasma and the brain during AD progression.

Extensive research describing the mechanisms underlying intraneuronal accumulation of Aβ peptides has been reported. Previous work conducted by us and others have demonstrated that Aβ40 and Aβ42 are internalized by distinct mechanisms^14–15^ and follow different itineraries in neuronal cells^16^. However, similar studies have not been conducted in BBB endothelial cells.

In the current study, we systematically dissected the cellular uptake and intracellular itineraries of fluorescently labeled soluble Aβ40 and Aβ42 (F-Aβ40 or F-Aβ42) in BBB cell culture models in vitro. We quantified the intracellular accumulation of F-Aβ using flow cytometry and confocal microscopy following pharmacological inhibition and siRNA knockdowns to investigate the contributions of known endocytic mechanisms. Further, we assessed the intracellular distributions of F-Aβ isoforms by labeling various endo-lysosomal organelles followed by live cell imaging. Our studies revealed that F-Aβ40 and F-Aβ42 are endocytosed by BBB endothelial cells via distinct molecular pathways and are differentially sorted by the endo-lysosomal system, which could lead to their distinct trafficking and accumulation kinetics in the BBB endothelium.

## RESULTS

### Energy dependent uptake of F-Aβ40 and F-Aβ42 in BBB endothelial cells

Receptor-mediated endocytosis requires energy produced by ATP hydrolysis. To determine if the uptake of F-Aβ40 or F-Aβ42 in human cerebral microvascular endothelial cell (hCMEC/D3) monolayers is mediated by energy-dependent endocytosis, the extent of F-Aβ internalization with or without ATP depletion was determined by flow cytometry **(Figure 1A)**. As shown in the histograms, the intra-endothelial F-Aβ40 **(Figure 1B)** and F-Aβ42 **(Figure 1C)** accumulation was significantly reduced in ATP depleted cells compared to control cells. The corresponding geometric mean of fluorescence intensity and coefficient of variance were displayed in **Figure 1D**.

**Figure 1.**
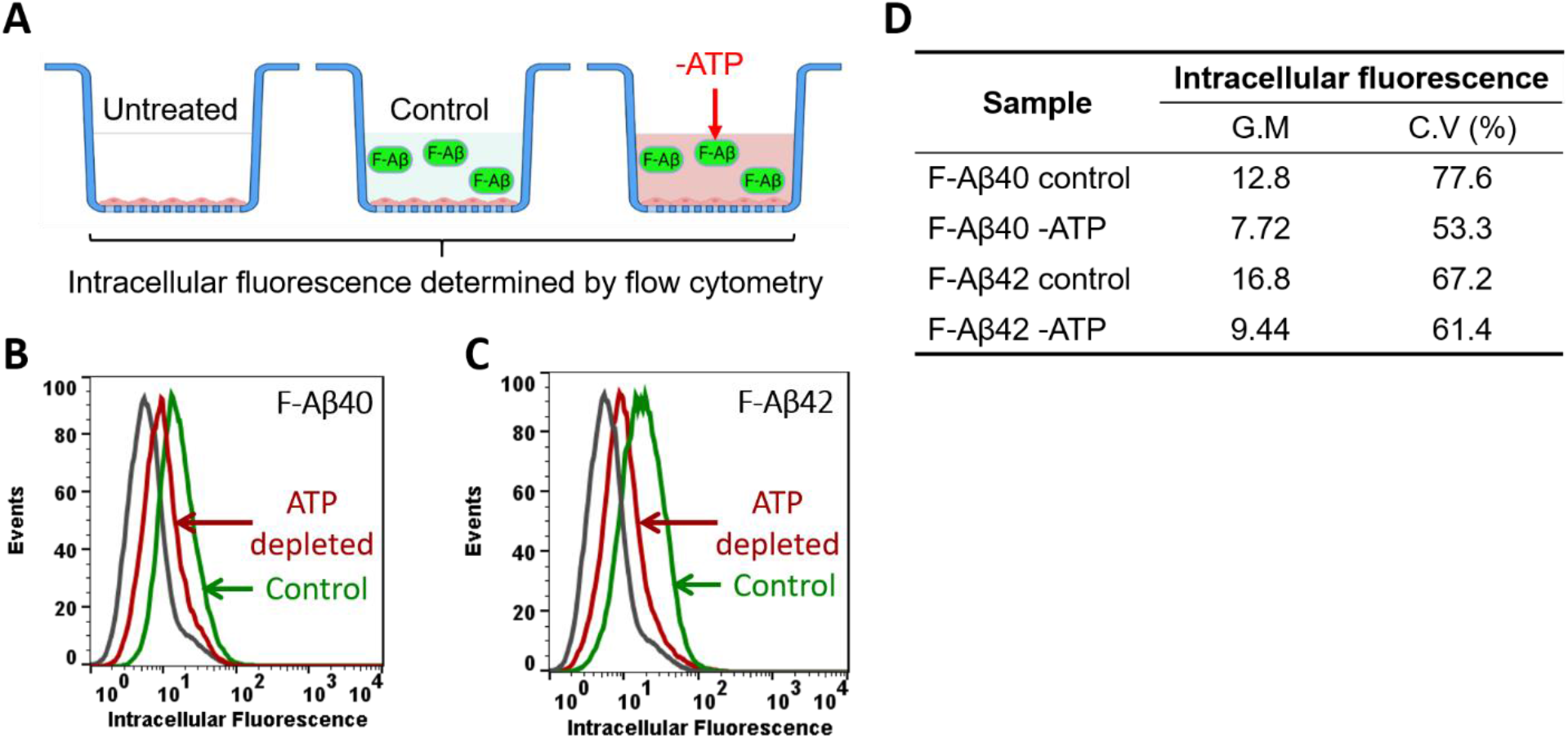
Energy dependent uptake of F-Aβ peptides in polarized hCMEC/D3 monolayers. **(A)** Experimental design. Cells grown on Transwell^®^ filters were pre-incubated with glucose free DMEM supplemented with 50mM 2-deoxy glucose and 0.1% sodium azide for 30 min. Then 1 μM F-Aβ was added to the luminal compartment and incubated for 30 min. Intracellular fluorescence was assessed by flow cytometry. **(B-C)** Histograms of hCMEC/D3 cells incubated with **(B)** F-Aβ40 or **(C)** F-Aβ42 in control and ATP depleted group. **(D)** Comparison of geometric means of F-Aβ40 and F-Aβ42 fluorescence in control and ATP depleted cells.

### Dynamin dependent uptake of F-Aβ40 and F-Aβ42 in BBB endothelial cells

To determine if F-Aβ endocytosis is dependent on dynamin, a protein involved in pinching off the endocytotic vesicles^17–18^, the effect of dynasore, a potent dynamin inhibitor, on F-Aβ uptake was assessed by flow cytometry **(Figure 2A)**. It was shown that uptake of F-Aβ40 was significantly inhibited in dynasore pre-treated cells compared to the control cells **(Figure 2B)**. Uptake of F-Aβ42 also appeared to decrease in dynamin treated cells compared to the control cells; however, the difference was not statistically significant **(Figure 2C)**. Effect of dynamin on F-Aβ internalization was further verified in hCMEC/D3 cells overexpressing dominant-negative mutant K44-dynamin (**Figure 2D**). Substantial decrease in green puncta of F-Aβ40 and red puncta of AF633-Trf were observed in cells expressing K44-dynamin mutant (**Figure 2F**) compared to control cells (**Figure 2E**). Similarly, uptake of F-Aβ42 was also inhibited in K44-dynamin transfected cells (**Figure 2H**) compared to control cells (**Figure 2G**). Moreover, cells that were transfected with GFP-K44 dynamin (green puncta, **Figure 2J**) clearly showed lower uptake of both SR101-Aβ40 (orange puncta, **Figure 2K**) and AF633-Trf (red puncta, **Figure 2L**). However, in the same image, non-transfected cells without GFP expression demonstrated the uptake of both SR101-Aβ40 and AF633-Trf.

**Figure 2.**
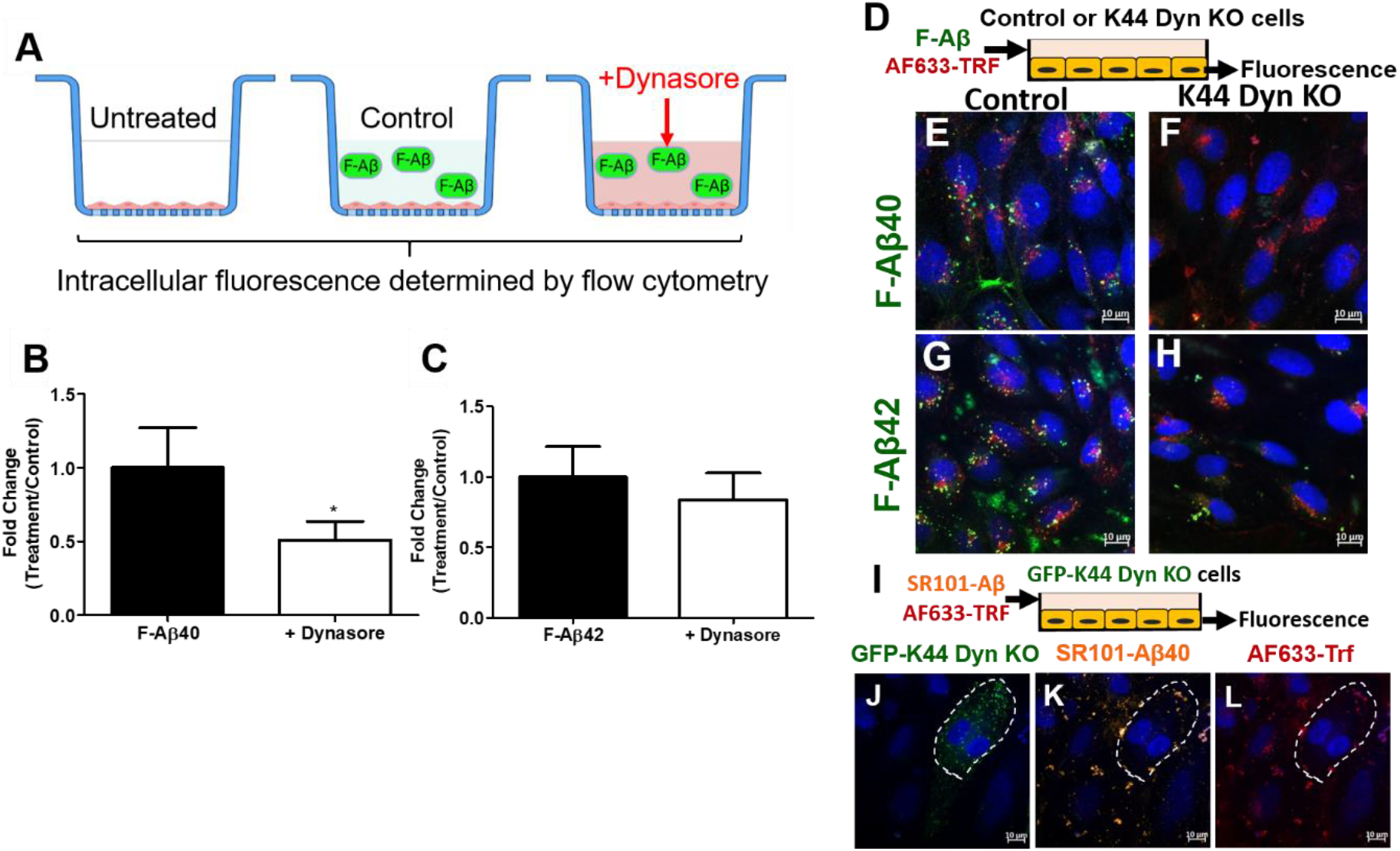
Dynamin dependent endocytosis of F-Aβ40 and F-Aβ42 in hCMEC/D3 cells. **(A)** Experiment design of flow cytometry study. Cells grown on Transwell^®^ filters were pre-incubated with 80 μM dynasore for 30 min. Then 1 μM F-Aβ was added to the luminal compartment and incubated for 60 min. Intracellular fluorescence was assessed by flow cytometry. **(B-C)** Fold change of median cellular fluorescence intensity from three replicates (n=3). *p<0.05, student’s t-test. **(D, I)** Experiment design of confocal microscopy. Cells grown on coverslip bottom dishes were transfected with GFP or non-GFP K44-negative-dominant dynamin. Transfected cells were incubated with 1 μM F-Aβ or Sulforhodamine101 labeled Aβ (SR-Aβ) for 60 min followed by 20 μg/ml of AlexaFluor labeled transferrin (AF633-TRF) treatment for 30 min. Following fixation and nuclear staining, the cells were imaged by laser confocal microscopy. **(E-H)** Confocal micrographs depicting internalization of F-Aβ (green) and AF633-Trf (red) in **(E, G)** control cells and in **(F, H)** cells expressing negative-dominant K44-dynamin mutant (K44 Dyn KO). **(J-L)** Intracellular uptake of AF633-Trf (red) and SR-Aβ40, orange) in cells expressing GFP labeled negative-dominant K44-Dynamin (GFP-K44 Dyn KO). **(J)** GFP-K44 Dyn KO cells marked by dashed white line, **(K)** SR-Aβ40, and **(L)** AF633-Trf.

### Actin dependent uptake of F-Aβ40 and F-Aβ42 in BBB endothelial cells

To investigate the impact of actin on F-Aβ uptake. The intracellular accumulation of F-Aβ40 and F-Aβ42 in hCMEC/D3 cells was determined by flow cytometry with or without actin inhibition **(Figure 3A)**. The histograms demonstrated that uptake of F-Aβ40 **(Figure 3B)** and F-Aβ42 **(Figure 3C)** was substantially reduced in cells pre-treated with cytochalasin A, which disrupts actin dynamics. Further, the reductions were found to be statistically significant for both F-Aβ40 **(Figure 3D)** and F-Aβ42 **(Figure 3E)** from three independent experiments.

**Figure 3.**
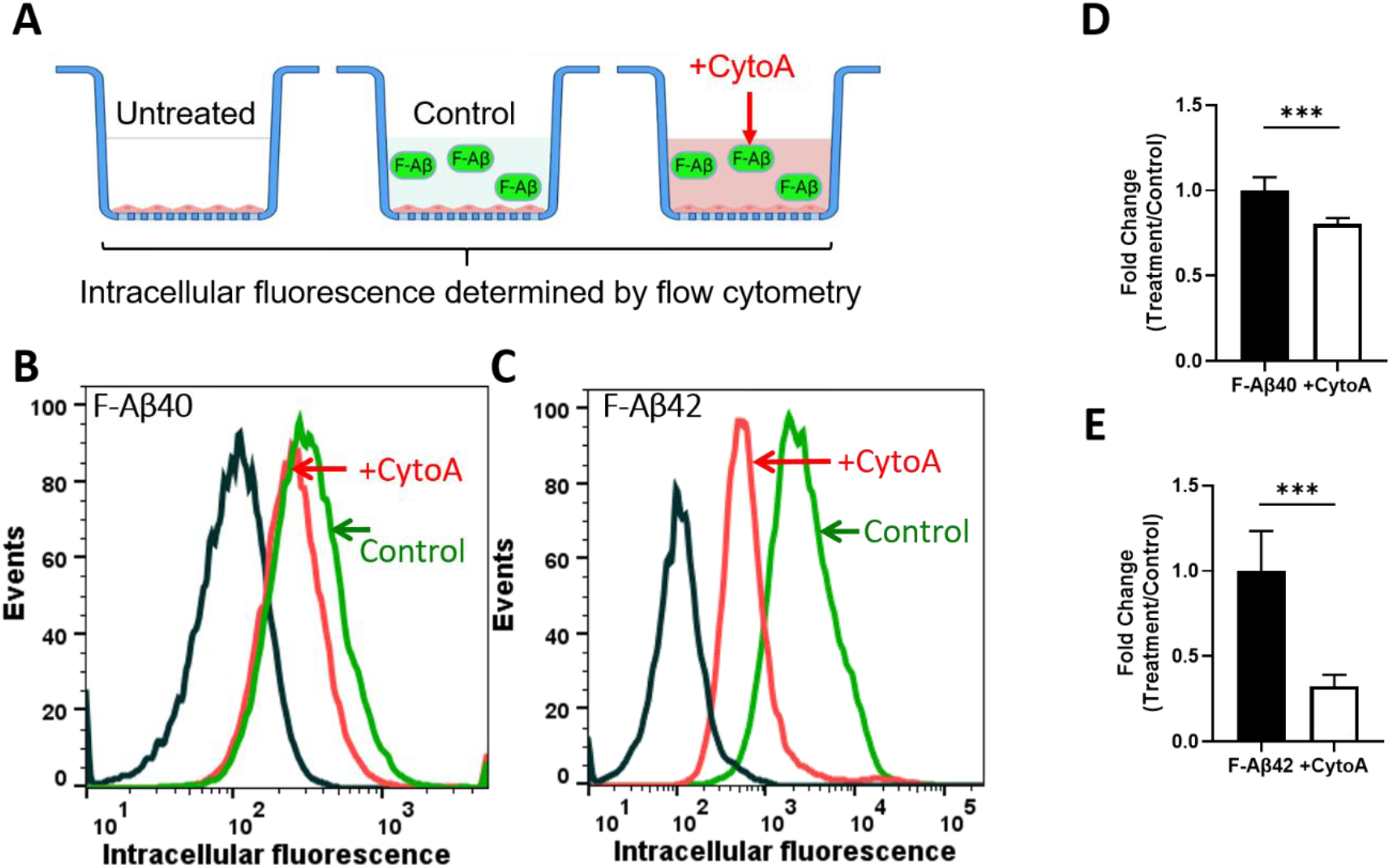
Actin dependent endocytosis of F-Aβ peptides in polarized hCMEC/D3 monolayers. **(A)** Experimental design. Cells were pre-incubated with cytochalasin A (CytoA,10 μM) for 30 min. Then 1 μM F-Aβ F-Aβ was added and incubated for 1 hour. Intracellular fluorescence was assessed by flow cytometry. **(B-C)** Histograms of hCMEC/D3 cells incubated with **(B)** F-Aβ40 or **(C)** F-Aβ42 in control and CytoA treated group. **(D-E)** Fold change of median cellular fluorescence intensity of the total number of gated cells for three replicate samples (n=3). ***p<0.001, student’s t-test.

### F-Aβ40 uptake by BBB endothelial cells via clathrin-mediated endocytosis

Flow cytometry analysis showed lower accumulation of F-Aβ40 in cells treated with monodansylcadaverine (MDC) (**Figure 4A**) and chlorpromazine (CPZ) (**Figure 4C**) and the difference was found to be statistically significant (**Figure 4E**). Although, F-Aβ42 uptake was reduced in MDC (**Figure 4B**) and CPZ (**Figure 4D**) pre-treated hCMEC/D3 cells, the difference was not significant (**Figure 4F**). To further confirm the role of clathrin-mediated endocytosis on F-Aβ uptake, knockdown of clathrin in hCMEC/D3 cells was performed using siRNA. Then the cells were incubated with AlexaFluor 633-transferrin (AF633-Trf) and F-Aβ **(Figure 4G)**. The AF633-Trf (red puncta, **Figure 4H-K**) was used as a marker of clathrin-mediated endocytosis and its uptake was inhibited in the cells transfected with clathrin siRNA compared to the control cells that underwent mock transfection. In these cells, intracellular uptake of F-Aβ40 was also significantly reduced compared to the control cells (green puncta, **Figure 4H-I**). However, no substantial change was observed in F-Aβ42 uptake (green puncta, **Figure 4J-K**).

**Figure 4.**
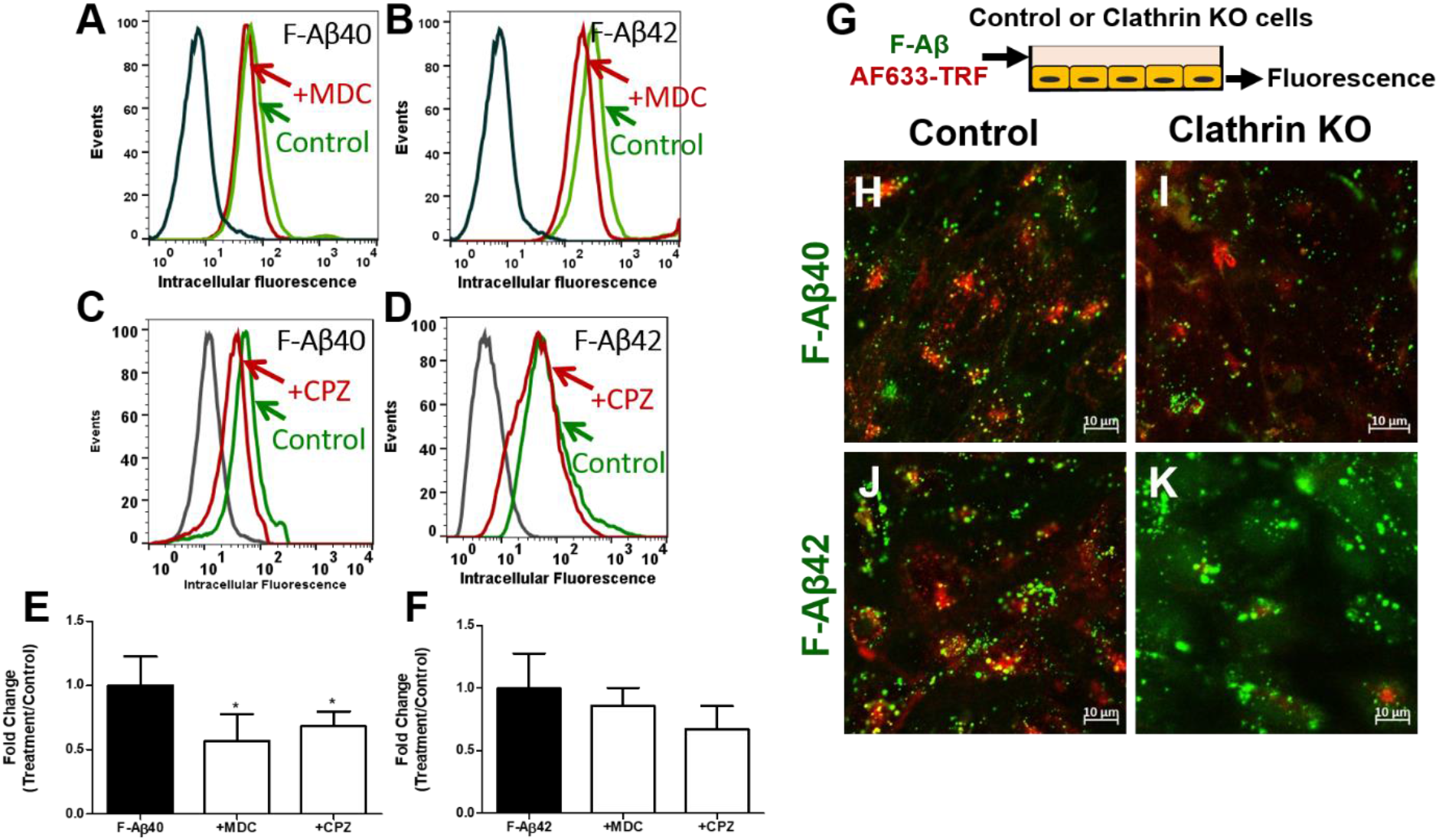
Role of clathrin-mediated endocytosis in the internalization of F-Aβ40 and F-Aβ42 in polarized hCMEC/D3 monolayers. **(A-D)** Histograms of hCMEC/D3 cells incubated with **(A)** F-Aβ40 and **(B)** F-Aβ42 in control and monodancyl cadverin (MDC, a specific clathrin inhibitor) pretreated group, **(C)** F-Aβ40 and **(D)** F-Aβ42 in control and chlorpromazine (CPZ, a non-specific clathrin inhibitor) pretreated group. **(E-F)** Fold change of median cellular fluorescence intensity from three replicates (n=3). *p<0.05 compared to F-Aβ alone group, one-way ANOVA with Bonferroni post-tests. **(G)** Experiment design of confocal microscopy study. Control or clathrin knocked-down cells were treated with 1 μM F-Aβ peptides and 20 μg/mL AlexaFluor 633-transferrin (AF633-Trf) and then subjected to confocal microscopy. **(H-K)** Confocal micrographs depicting internalization of F-Aβ (green) and AF633-Trf (red) in **(H, J)** control cells and **(I, K)** clathrin knocked down cells.

### F-Aβ42 uptake by BBB endothelial cells via caveolae-mediated endocytosis

Flow cytometry studies conducted on hCMEC/D3 cells demonstrated that nystatin (**Figure 5B**) or methyl-β-cyclodextrin (mβCD) (**Figure 5D**) treatment significantly reduced the F-Aβ42 uptake (**Figure 5F)**. However, no impact of nystatin on F-Aβ40 uptake was observed as indicated by the histogram (**Figure 5A**) and statistical analysis (**Figure 5E**), although the fluorescence intensity of F-Aβ40 was still significantly reduced after mβCD treatment (**Figure 5C, E**). In a separate study, primary bovine brain microvascular endothelial (BBME) cells grown on Transwell^®^ inserts were treated with mβCD and F-Aβ, followed by confocal microscopy imaging (**Figure 5G**). The z-stack images demonstrated reduced intracellular uptake of F-Aβ42 (green puncta) in mβCD treated monolayers (**Figure 5K**) in comparison to control cells (**Figure 5J**). In contrast, the uptake of F-Aβ40 was unaffected by mβCD pretreatment (**Figure 5H-I**).

**Figure 5.**
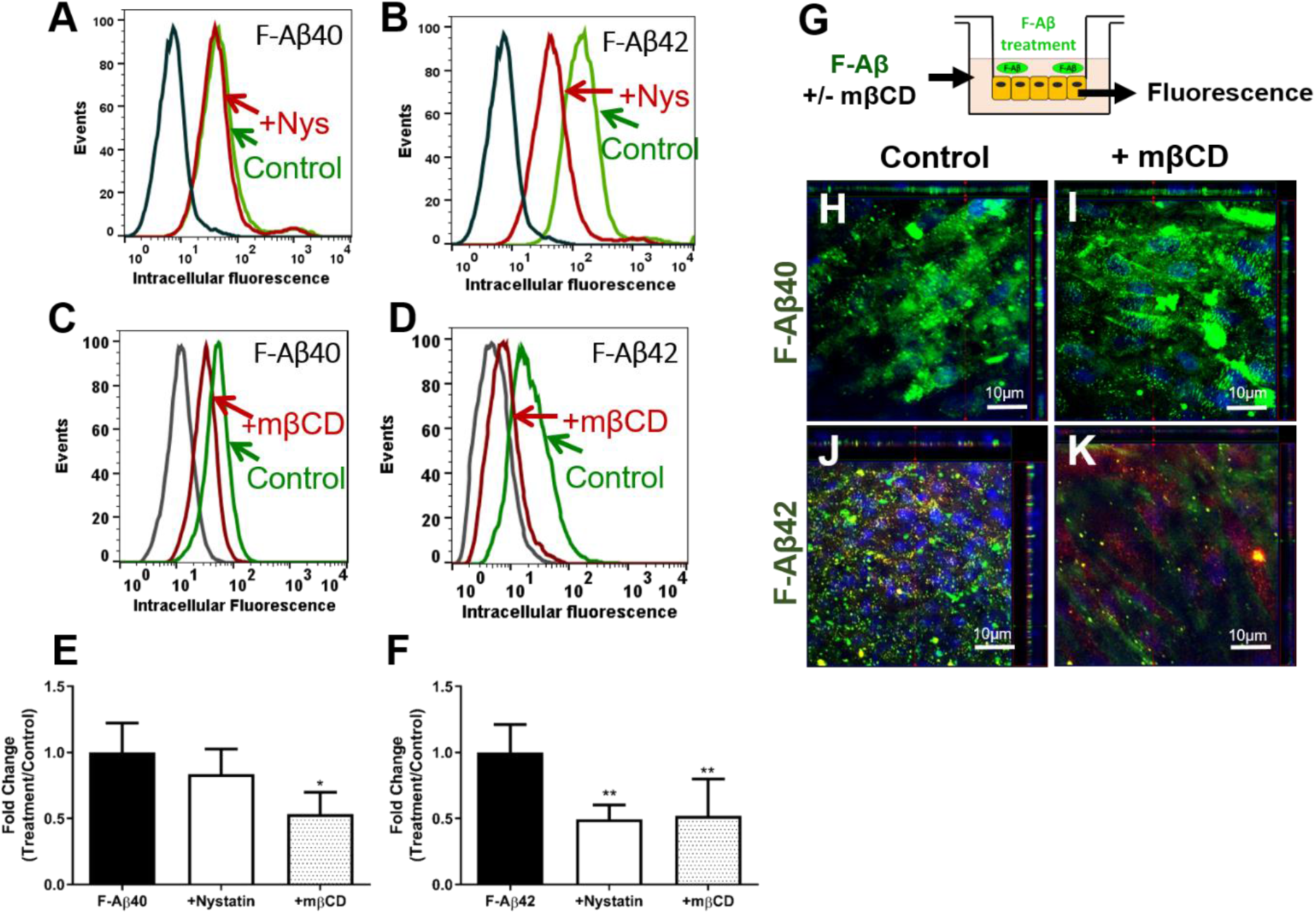
Role of lipid rafts on the internalization of F-Aβ peptides in hCMEC/D3 and bovine brain microvascular endothelial (BBME) cells. **(A-D)** Histograms of hCMEC/D3 cells incubated with **(A)** F-Aβ40 and **(B)** F-Aβ42 in control and Nystatin (a specific inhibitor of lipid raft-caveolae endocytosis pathway) pretreated group, **(C)** F-Aβ40 and **(D)** F-Aβ42 in control and mβCD (a non-specific cholesterol chelator) pretreated group. **(E-F)** Fold change of median cellular fluorescence intensity Fold change of median cellular fluorescence intensity from three replicate samples (n=3). *p<0.05, **p<0.01 compared to F-Aβ alone group, one-way ANOVA with Bonferroni post-tests **(G)** Experiment design of confocal microscopy study. Polarized BBME cell monolayers cultured on Transwell® filters were preincubated with or without methyl beta cyclodextrin (mβCD) in DMEM and then incubated with 1 μM F-Aβ peptides and/or 20 μg/ml AF633-Trf. **(H-K)** The z-series confocal micrographs demonstrating F-Aβ (green) fluorescence in **(H, J)** control cells and **(I, K)** mβCD treated cells. Images were shown in x-y (large square), x-z (top horizontal panel) and y-z (right vertical panel) plane.

### Tyrosine phosphorylation of caveolin-1 is induced by Aβ42 but not by Aβ40 and mediates Aβ42 uptake

Caveolin-1 is a major structural component of caveolae, to further confirm the impact of caveolin-1 on cellular uptake of Aβ, knockdown of caveolin-1 in hCMEC/D3 cells was performed using siRNA and intracellular Aβ was measured by ELISA. Successful knockdown of caveolin-1 was demonstrated by western blotting as shown in **Figure 6A**. Moreover, caveolin-1 knockdown was shown to significantly reduce Aβ42 cellular uptake whereas no significant difference was observed for Aβ40 (**Figure 6B)**. It has been reported that phosphorylation of caveolin-1 is associated with caveolae-dependent endocytosis^19^‘^20^. In order to study the effect of Aβ40 and Aβ42 on the phosphorylation of caveolin-1, we exposed hCMEC/D3 cells to soluble Aβ peptides predominantly containing monomers and determined the phospho-caveolin-1 expression by western blotting. Incubation with Aβ42 resulted in a significant increase in the phosphorylation of caveolin-1 on tyrosine 14 and pretreatment of Src kinase inhibitor PP1 was shown to block the phosphorylation (**Figure 6C**). By contrast, Aβ40 has no impact on caveolin-1 phosphorylation (**Figure 6D**).

**Figure 6.**
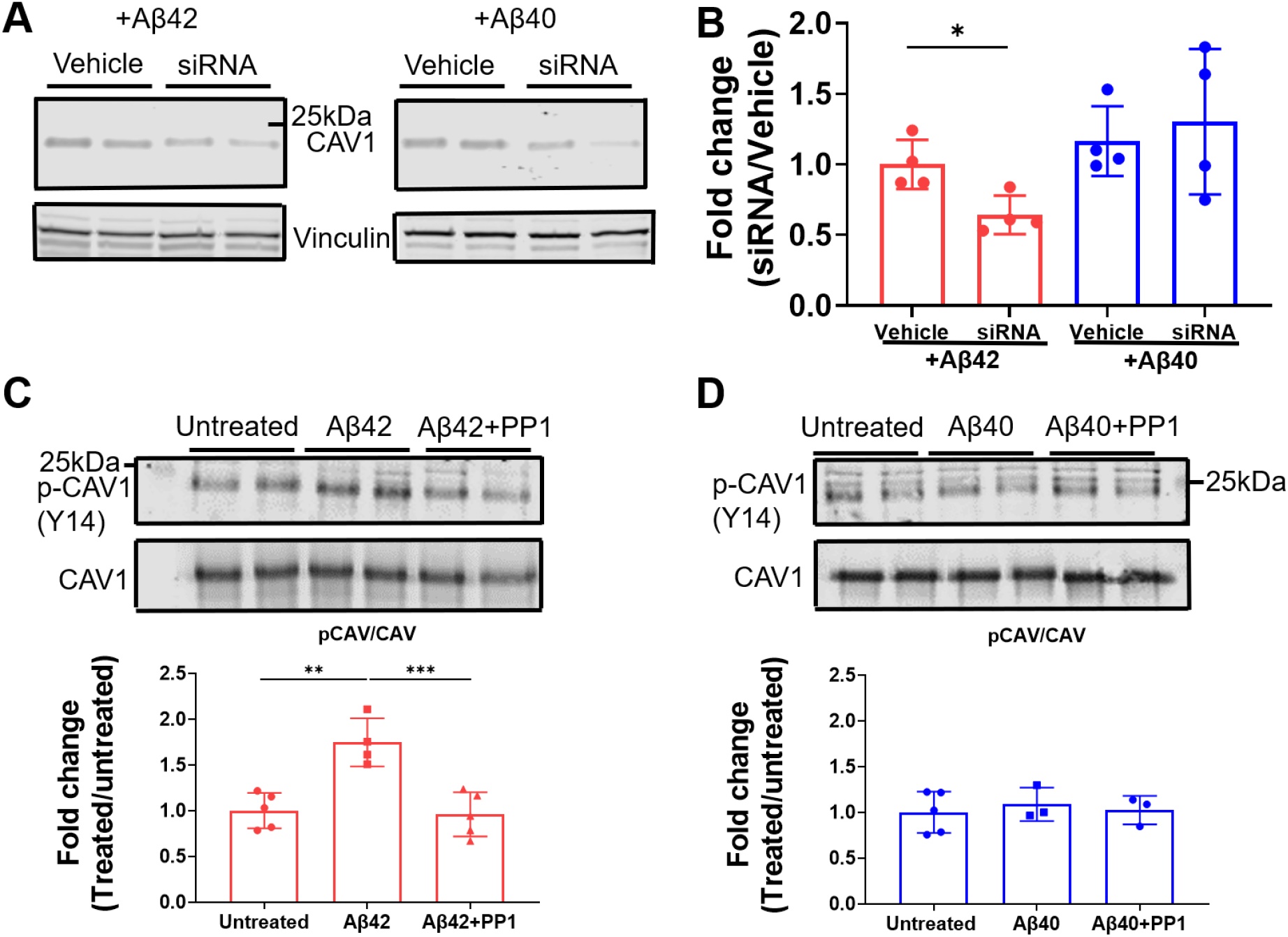
Tyrosine phosphorylation of caveolin-1 is induced by Aβ42 but not by Aβ40 and mediates Aβ42 uptake. **(A)** Representative immunoblots confirming caveolin-1 knockdown using siRNA in hCMEC/D3 cells. **(B)** Intracellular Aβ accumulation in hCMEC/D3 cells transfected with caveolin-1 siRNA. *p<0.05, student’s t-test. **(C-D)** Representative immunoblots and quantification of phosphorylated caveolin-1 (p-CAV1 (Y14)) and total caveolin-1 (CAV1) with **(C)** Aβ42 and **(D)** Aβ40 stimulation. ** p<0.01, ***p<0.005, one-way ANOVA with Boferroni post-tests.

### Intracellular itineraries of Aβ40 versus Aβ42 in BBB endothelial cells

#### Accumulation of Aβ40 and Aβ42 in early and late endosomes

hCMEC/D3 cells were transfected with m-cherry Rab5 protein (marker for early endosome) and then subjected to incubation with F-Aβ40. Confocal microscopy studies demonstrated that Aβ40 accumulated in the early endosomes as shown by substantial colocalization of F-Aβ40 (**Figure 7B**) and m-cherry Rab5 (**Figure 7A**). Further, F-Aβ40 was found to accumulate in the late endosomes, as indicated by colocalization of F-Aβ40 and Dil-LDL (**Figure 7D-F**). To confirm the accumulation of F-Aβ in the late endosomes, hCMEC/D3 cells transfected with GFP-labeled Rab7 protein (marker for late endosome) were incubated with SR-Aβ40 or SR-Aβ42 and then imaged live for up to 60 minutes. Only minor accumulation of SR-Aβ40 in the late endosomes was observed at early time points (**Figure 8A-B**). Contrarily, substantial colocalization between SR-Aβ42 and GFP-labeled Rab7 was found after 14 minutes of incubation (**Figure 8D-E**). Both peptides were shown to accumulate in the late endosome at later time points (**Figure 8C, F**).

**Figure 7.**
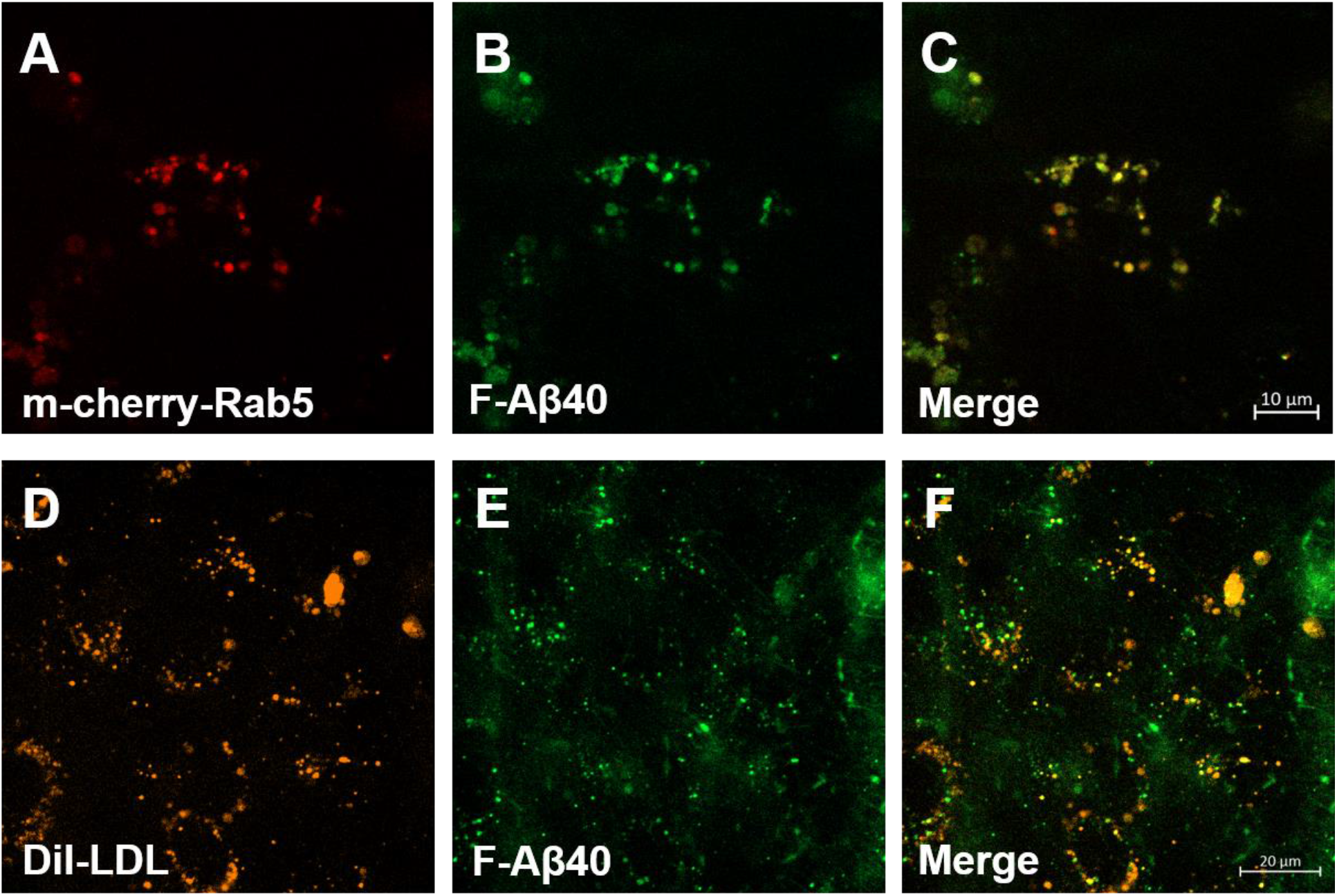
Accumulation of F-Aβ40 peptides in early and secondary endosomes in hCMEC/D3 cells. **(A-C)** Colocalization of F-Aβ40 with m-cherry fluorescent Rab5. hCMEC/D3 cells were transfected with m-cherry fluorescent Rab5 and then incubated with F-Aβ40 for 15 minutes. Then Live cell imaging was conducted after washing the cells with DMEM. **(A)** m-cherry fluorescent Rab5, **(B)** F-Aβ40, **(C)** Merge image of A and B. **(D-F)** Colocalization of F-Aβ40 with Dil-LDL. hCMEC/D3 cells were treated with 1 μM F-Aβ40 and Dil-LDL (15 μg/ml) for 60 minutes. Then Live cell imaging was conducted after washing the cells with DMEM. **(D)** Dil-LDL, **(E)** F-Aβ40, **(F)** Merge image of D and E.

**Figure 8.**
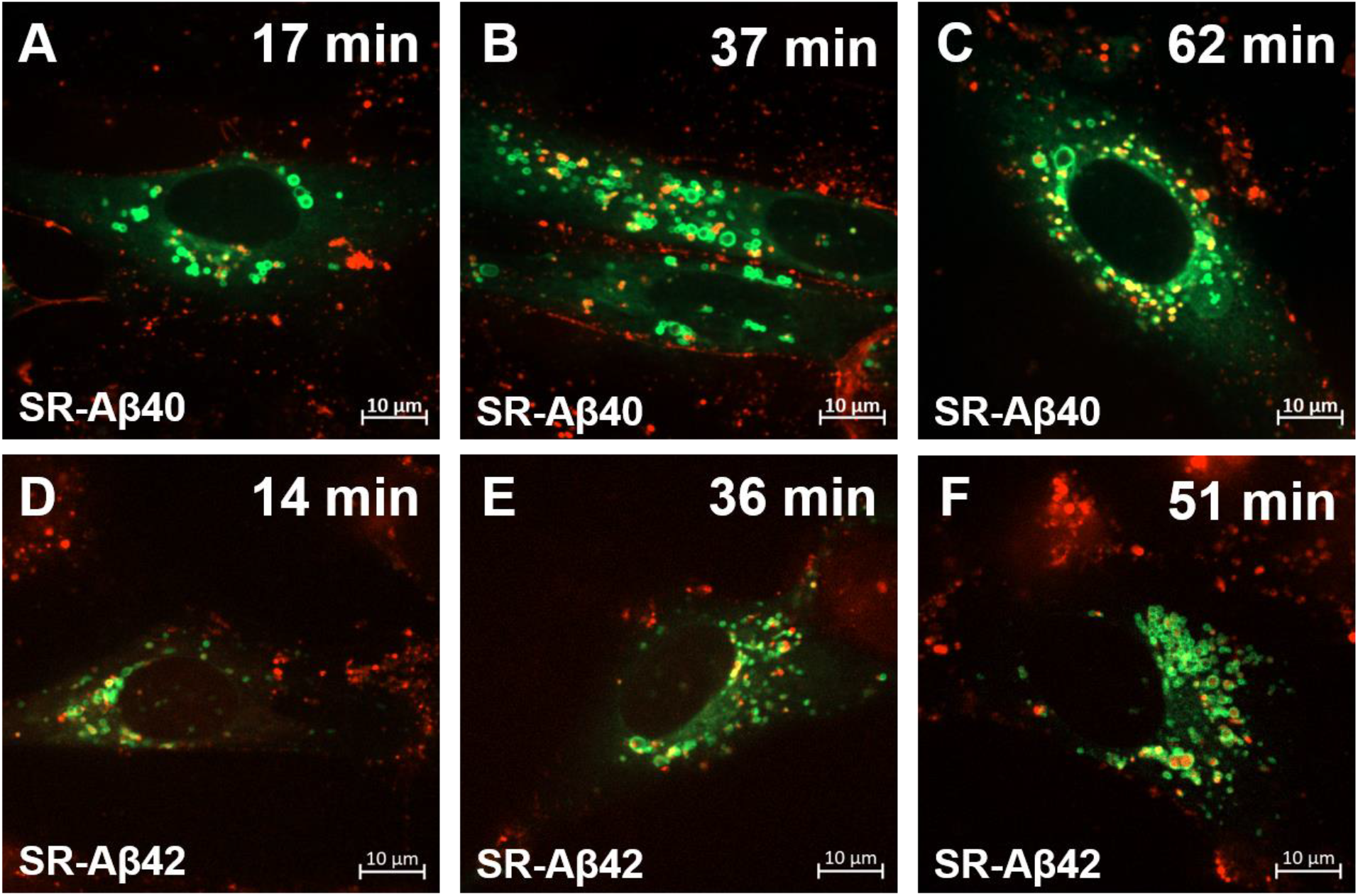
Time dependent accumulation of sulforhodamine labeled Aβ (SR-Aβ) peptides in late endosomes. hCMEC/D3 cells expressing GFP-Rab7 incubated with SR-Aβ for 15 minutes. Then the cells were washed with DMEM and imaged by live cell imaging up to 60 minutes. **(A-C)** Accumulation of SR-Aβ40 following incubations at **(A)** 17min **(B)** 37 min **(C)** 62 min; **(D-E)** Accumulation of SR-Aβ42 following incubations at **(D)** 14min **(E)** 36 min **(F)** 51 min.

#### Accumulation of Aβ40 in recycling endosomes

Accumulation of Aβ40 in recycling endosomes was examined in hCMEC/D3 cells transfected with GFP-labeled Rab11, which is a marker for recycling endosomes. Considerable colocalization of SR-Aβ40 and GFP-labeled Rab11 was observed throughout the incubation period (**Figure 9**), confirming Aβ40 accumulation in recycling endosomes.

**Figure 9.**
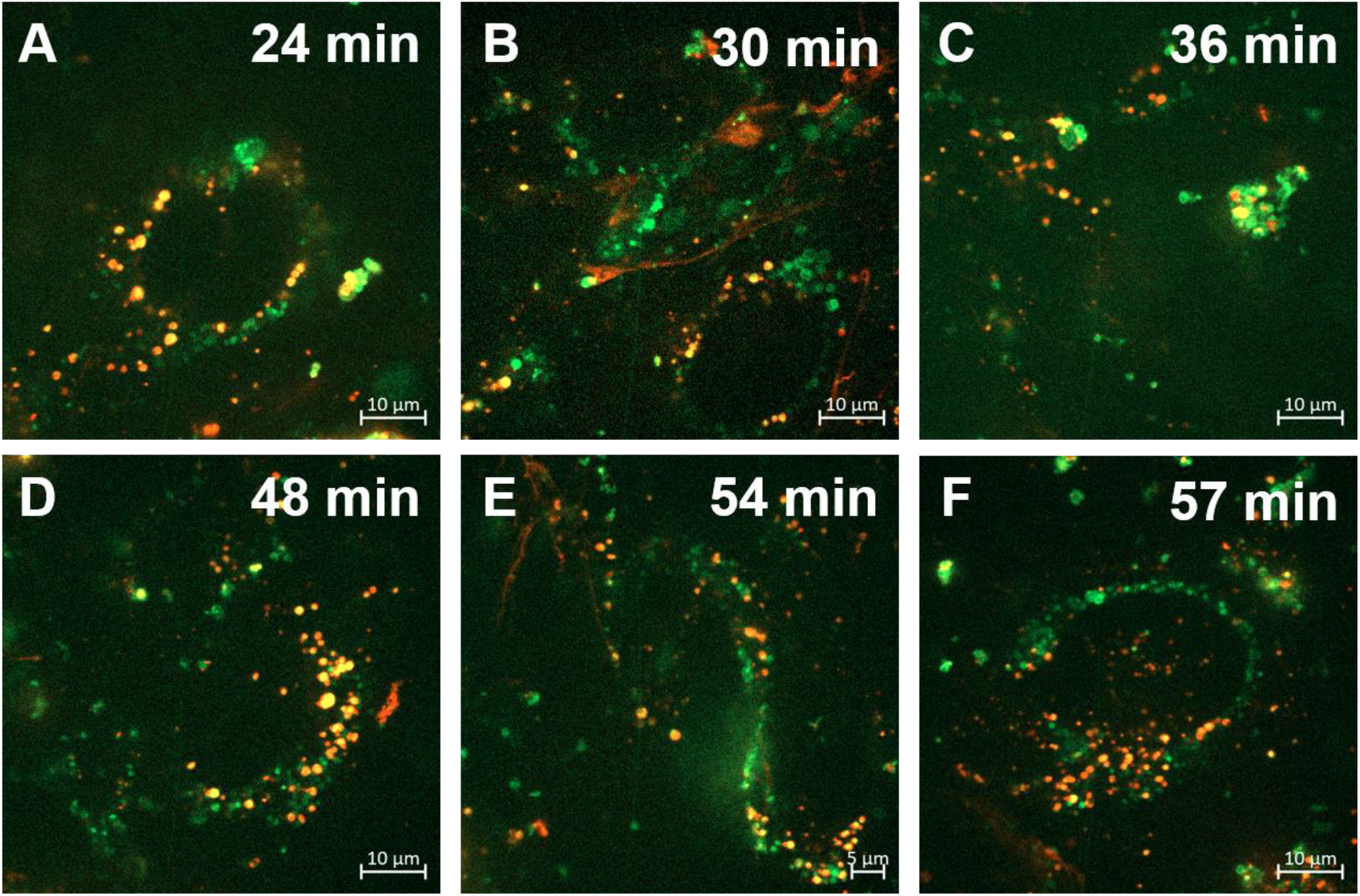
Time dependent accumulation of sulforhodamine labeled Aβ40 peptides in recycling endosomes. hCMEC/D3 cells expressing GFP-Rab11 incubated with SR-Aβ40 for 15 minutes. Then the cells were washed with DMEM and imaged by live cell imaging up to 60 minutes. **(A-C)** Accumulation of SR-Aβ40 following incubations at **(A)** 24min **(B)** 30 min **(C)** 36 min **(D)** 48min **(E)** 54 min **(F)** 57 min.

#### Accumulation of Aβ40 and Aβ42 in lysosomes

hCMEC/D3 cells were co-incubated with F-Aβ and lysotracker (predominantly a marker for lysosomes) to examine lysosomal accumulation of F-Aβ. The F-Aβ40 displayed only a partial localization in the lysosomes (Figure 10A-D), whereas F-Aβ42 exhibited considerable lysosomal accumulation (Figure 10E-H). Pearson’s correlation coefficient of F-Aβ42, which describes its extent of colocalization with lysosomes was found to be significantly higher than that of F-Aβ40 (**Figure 10I**).

**Figure 10.**
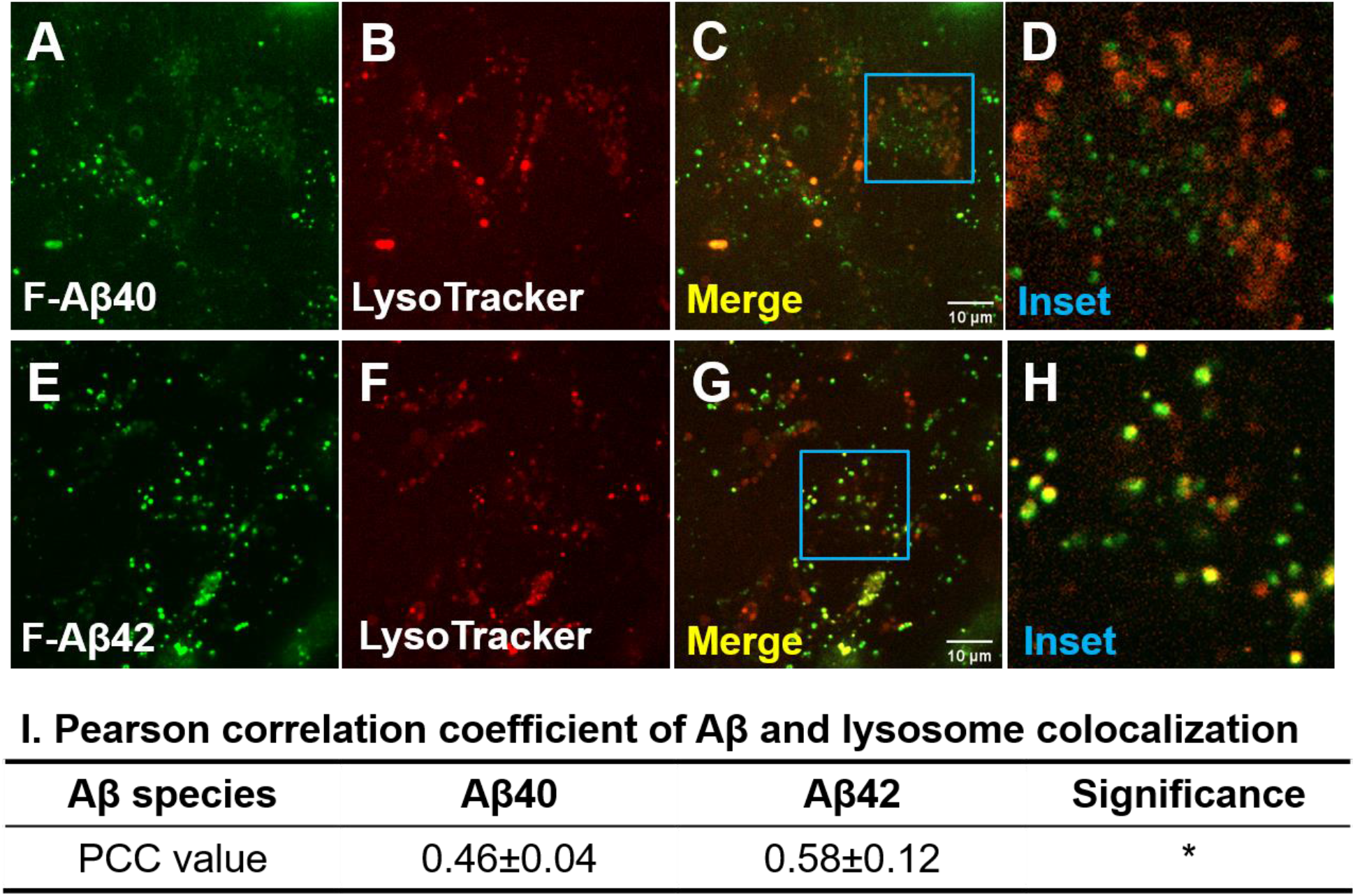
Accumulation of F-Aβ40 and F-Aβ42 peptides in lysosomes in hCMEC/D3 cells. Representative images showing colocalization of F-Aβ (green) and LysoTracker^™^ (red) in hCMEC/D3 cells. **(A-C)** F-Aβ40; **(E-G)** F-Aβ42; **(D, H)** Inset showing the magnified images of C & G. (I) Pearson correlation coefficient of F-Aβ and LysoTracker^™^. *p<0.05, student’s t-test

## DISCUSSION

BBB is a pivotal portal that regulates the bi-directional transport of Aβ mostly via receptor-mediated transcytosis, involving endocytosis, intracellular sorting, and exocytosis to the contralateral side^21^‘^22^. Multiple receptors such as RAGE and LRP1 have been proposed to regulate influx and efflux of Aβ, respectively. Although both Aβ40 and Aβ42 were reported to be substrates of these receptors/transporters, these peptides exhibited distinct disposition profiles, and are differently impacted by various pathophysiological conditions. It has been previously reported that Aβ40 is more vasculotropic than Aβ42 and primarily accumulate as amyloid deposits in the brain blood vessels, whereas Aβ42 forms the core of amyloid plaques that accumulate in the brain parenchyma^23^. Moreover, the CSF levels of soluble Aβ40 and Aβ42 decrease with aging in AD transgenic mice^24^ and AD patients^2^. However, the magnitude of Aβ42 decrease is greater than that of Aβ40, thus resulting in a reduction in Aβ42/Aβ40 ratio. Therefore, Aβ42/Aβ40 ratio in CSF rather than Aβ42 concentration alone is considered as a reliable biomarker to detect AD. Hence investigating molecular mechanisms that disrupt Aβ42/Aβ40 ratio during AD progression is expected to inform key pathophysiological pathways driving AD progression.

Although, several investigators opined that the reduction in soluble Aβ42/Aβ40 ratio is due to the propensity of Aβ42 to form insoluble fibril and amyloid plaques, contribution of Aβ42 and Aβ40 trafficking disruptions at the BBB endothelium that could affect the levels of both isoforms in CSF and plasma cannot be ruled out. Bioinformatic analysis has shown that the genes encoding for various proteins involved in intracellular cargo sorting are downregulated in brains of AD patient brains^25^. Therefore, we hypothesize that the BBB endothelium discriminates Aβ isoforms and traffics them independently via distinct transcytosis mechanisms. Selective disruption of transcytosis pathways that predominantly handle Aβ40 versus Aβ42 at the BBB may contribute to variations in Aβ42/40 ratios observed during AD progression.

Moreover, dysregulated trafficking of Aβ peptides could promote their accumulation in the BBB endothelium and induce BBB dysfunction, which is associated with AD progression. However, the molecular mechanisms regulating intracellular trafficking and accumulation of Aβ in the BBB endothelial cells remain elusive. Therefore, in this study, we characterized the cellular mechanisms involved in Aβ internalization and intracellular transit at the BBB endothelium.

Our previous studies have shown that the uptake of Aβ40 by bovine brain microvascular endothelial cells is temperature dependent^14^. Current study confirmed that the uptake of both Aβ40 and Aβ42 in hCMEC/D3 monolayers require energy and is inhibited upon ATP depletion (**Figure 1**). Further, our recent publication showed saturable uptake of Aβ by BBB endothelium^26^. Taken together, these results demonstrated that both Aβ40 and Aβ42 are internalized by BBB endothelial cells via receptor-mediated endocytosis. Next, we have shown that Aβ internalization is dynamin dependent. Dynamin is recruited to the necks of vesicles formed during endocytosis and induces membrane scission to release vesicles into the cytosol^17^‘^18^. By pretreating hCMEC/D3 monolayers with dynasore (dynamin inhibitor) followed by flow cytometry and transfection of mutant form of dynamin followed by confocal microscopy, we demonstrated that dynamin is involved in the endocytosis of both F-Aβ40 and F-Aβ40 (**Figure 2**). Previous studies have shown that the uptake of Aβ42 but not Aβ40 is reduced in dynasore treated PC-12 cells^16^ and SH-SY5Y cells^15^. The discrepancy between BBB endothelial cells and neuronal cells could be due to differential expression of dynamin isoforms. While dynamin 1 and 3 are highly expressed in neurons, endothelial cells are believed to mostly express dynamin 2^27^.

During endocytosis, actin polymerization provides the force for vesicle budding and has been reported to mediate uptake of cargos in various cells^28^‘^29^. In the current study, we found that both Aβ40 and Aβ42 utilize actin-dependent endocytic mechanisms to enter the BBB endothelial cells (**Figure 3**). Similar results were reported in neurons^15^ and astrocytes^30^, where disruption of actin polymerization by cytochalasin D was shown to significantly reduce Aβ40 and Aβ42 endocytosis. These studies have indicated that both Aβ40 and Aβ42 are internalized by BBB endothelial cells via energy, dynamin and actin-dependent endocytosis.

We further investigated specific mechanisms involved in Aβ endocytosis. Clathrin and caveolae-mediated endocytosis are the two well-studied mechanisms and were shown to facilitate the receptor-mediated transcytosis (RMT) of various macromolecules at the BBB. Endocytosis of critical endogenous proteins such as transferrin^31^, LDL^32^, and insulin^33^ is mediated by clathrin. In contrast, caveolae-mediated transcytosis mediates the blood-to-brain entry of viruses and extracellular proteins such as albumin^34^. Dysregulated caveolae-mediated transcytosis is believed to increase BBB permeability under pathological conditions triggered by ischemic stroke^35^ and sub-arachnoid hemorrhage^36^ or under exposure to low-intensity focused ultrasound^37^. Therefore, we investigated the involvement of both clathrin and caveolae mediated endocytic mechanisms in Aβ uptake at the BBB endothelium. Using chemical inhibitors that inhibit the clathrin-coated pit assembly at the plasma membranes, MDC and CPZ^38–39^, we have shown that the endocytosis of Aβ40 but not Aβ42 is clathin-mediated in hCMEC/D3 monolayers (**Figure 4A-F**). These results were further verified by confocal microscopy which demonstrated significant reduction in the uptake of F-Aβ40 but not F-Aβ42 by hCMEC/D3 monolayers in which clathrin was knockdown using siRNA (**Figure 4H-K**). It has been reported that clathrin-mediaed endocytosis pathway is disrupted in AD^40–42^, which is also reflected in impaired Aβ40 endocytosis at the BBB. Indeed, Zhao et. al have found that PICALM regulates clathrin-dependent internalization of Aβ bound to LRP1, leading to BBB-mediated clearance of Aβ. Further, PICALM levels and Aβ clearance were shown to be reduced in AD-derived endothelial monolayers^43^. On the other hand, inhibitors of caveolae-mediated endocytosis were shown to significantly decrease the uptake of Aβ42 but not Aβ40 as shown by confocal microscopy (**Figure 5H-K**). Although mβCD also significantly reduced the Aβ40 uptake in flow cytometry analysis, this inhibitor is non-specific and acute cholesterol depletion by mβCD has also been reported to inhibit clathrin-mediated endocytosis^44^. Involvement of caveolae-mediated endocytosis in Aβ42 uptake was further confirmed by siRNA knockdown studies showing that caveolin-1 knockdown significantly reduced Aβ42 uptake but had no impact on Aβ40. Moreover, western blot results demonstrated that only Aβ42, not Aβ40, increased the tyrosine-14 phosphorylation of caveolin-1 (**Figure 6**). Src-dependent caveolin-1 phosphorylation is suggested to facilitate the formation and internalization of caveolae into the cytoplasm^19, 45-46^; phosphorylation associated conformational change of caveolin-1 is believed to engender maturation of caveolae and subsequent release from the plasma membrane^20^. Together, these studies indicate that the endocytosis of Aβ40 at the BBB endothelium is primarily clathrin-dependent whereas the endocytosis of Aβ42 is caveolae-mediated.

Differential involvement of endocytic pathways between Aβ40 and Aβ42 could provide mechanistic foundations underlying differential transport kinetics of these two peptides across the BBB in various pathophysiological conditions. For example, our previous study illustrated that peripheral insulin exposure decreased the brain influx of Aβ42 but increased Aβ40 influx. Further, insulin treatment in vitro enhanced the uptake of transferrin, a clathrin-mediated endocytosis marker, by BBB endothelial cells but reduced caveolae-mediated endocytosis of AF647-cholera toxin-B^47^. Hence, the differential effects of insulin on Aβ trafficking could be due to distinct endocytic pathways employed by these two peptides, which allows insulin to activate one while inhibiting the other. The molecular mechanism by which insulin exerts such selective effect on clathrin and caveolae-mediated endocytosis is not fully understood and the role of insulin signaling pathway in mediating these processes is currently being investigated in our lab. In another study, we demonstrated that blood-to-brain influx of Aβ40 decreased whereas Aβ42 increased with aging in WT mice^48^. Age-related shift from ligand-specific transport to non-specific caveolar transcytosis at the BBB has been previously reported^49^. Given the current finding that Aβ42 endocytosis at the BBB endothelium is caveolae-mediated, preferential influx of Aβ42 during aging is most likely due to the shifts in transcytosis that involves clathrin versus caveolae-coated vesicles.

Following the receptor-mediated endocytosis, cargos are sorted via the endo-lysosomal system constituting of early, late, recycling endosomes and lysosomes. In polarized BBB endothelial monolayers, such sorting system determines the fate of lysosomal degradation, recycling back to the plasma membrane, or transcytosis^50–52^. Therefore, in the second part of this study, we investigated the intracellular itineraries of Aβ40 and Aβ42 in polarized hCMEC/D3 cell monolayers. We found that the majority of internalized Aβ40 trafficked through the early and late endosomes as assessed by the co-localization of F-Aβ40 with m-cherry-Rab5 and Dil-LDL (**Figure 7**). Accumulation of Aβ40 and Aβ42 in the early endosomes was also reported in differentiated PC12 cells^16^ and mouse neuroblastoma N2a cells^53^. From the early endosome, Aβ is recycled to the plasma membrane or emptied into late endosomes and lysosomes for degradation. Live cell imaging has shown that SR-Aβ40 accumulation in late endosomes occurred with longer incubation times (40-60 minutes). In contrast, SR-Aβ42 rapidly entered late endosomes with shorter as well as longer periods of incubation (**Figure 8**). Subsequently, greater accumulation of F-Aβ42 was observed in the lysosomal compartment compared to F-Aβ40 (**Figure 10**). These results were consistent with our previous findings in neuronal cells^16^ and implies that Aβ42 is more susceptible to lysosomal degradation compared to F-Aβ40. On the other hand, F-Aβ40 accumulated in GFP-Rab11 labeled recycling endosomes up to 1 hour of incubation (**Figure 9)**. The Rab11 has been reported to regulate exocytosis of recycling vesicles at the plasma membrane and a recent study suggested that it also mediates the transcytosis of polymeric immunoglobulin A (pIgA) across the polarized epithelial cells^54^‘^55^. Our previous study also demonstrated that the exocytosis rate of intracellular Aβ40 to the abluminal side (transcytosis) was significantly higher than that to the luminal side (recycling)^47^. These results indicate that internalized Aβ40 and Aβ42 follow distinct trafficking paths in the BBB endothelium, which may determine their transcytosis potential versus intracellular degradation.

Published studies indicated that caveolae-mediated vesicle trafficking directs monocarboxylic acid transporter 1 (Mct1) into late endosome/lysosome compartments in BBB endothelial cells, whereas clathrin-mediated endocytosis directs Mct1 to Rab11-positive recycling endosomes^56^. Therefore, caveolae-mediated endocytosis could sort Aβ42 for lysosomal degradation while clathrin-mediated endocytosis moves Aβ40 into recycling endosomes. Another potential reason for the selectivity between lysosomes and recycling endosomes could be related to their binding affinity to the receptors. Transferrin receptor antibodies are being employed as Trojan horses for delivering cargo across the BBB. Preclinical studies have shown that the transcytosis efficiency of this antibody is influenced by its affinity to the transferrin receptor. For instance, Johnsen et al. found that antibodies with low affinities mediated higher uptake of gold nanoparticles into the brain than the antibodies with higher affinities^57^. Both in vivo pharmacokinetic modeling^13^ and cellular transcytosis^26^ studies in our previous publications demonstrated that Aβ42 has lower Michaelis constant (Km) than Aβ40, suggesting higher binding affinity of Aβ42 to the BBB endothelium. Like observed with transferrin receptor antibody, it is likely that such high binding affinity reduces Aβ42 transcytosis and increases its likelihood of lysosomal entrapment.

Extensive accumulation of Aβ42 (from 25 nM in extracellular fluid to 2.5 μM in the lysosomal compartment) in the acidic environment of lysosomes could trigger its misfolding and enhance the formation of insoluble aggregates ^58^. The misfolded Aβ42 aggregates were shown to disrupt the integrity of endosomal-lysosomal system and decrease the uptake of ovalbumin in primary neurons^59^. It is possible that the accumulation of Aβ42 in the lysosomes of BBB endothelial cells could have similar detrimental effects and cause disrupted trafficking of various cargos via the BBB. Moreover, Aβ42 oligomers were found to damage tight junctional proteins and result in paracellular leakage via activation of autophagy and up-regulation of metalloproteinases in murine brain capillary endothelial cells (bEnd.3)^10, 60^.

Taken together, our study demonstrated that Aβ40 and Aβ42 are internalized by the BBB endothelial cells via dynamin and actin-dependent endocytosis but employ different endocytotic pathways. Endocytosis of Aβ40 was found to be clathrin-mediated whereas Aβ42 endocytosis is caveolae-mediated. Following endocytosis, both isoforms were sorted by the endo-lysosomal system; however, Aβ42 was shown to accumulate more in lysosomes than Aβ42, which could lead to higher degradation and/or aggregation of Aβ42. These findings were summarized in Figure 11. Our results provided molecular insights into mechanisms that regulate the transport Aβ40 versus Aβ42 in the BBB endothelium. This knowledge is critical to understand mechanisms underlying Aβ accumulation in the BBB endothelium and resultant BBB dysfunction, which is widely believed to aggravate AD progression.

**Figure 11.**
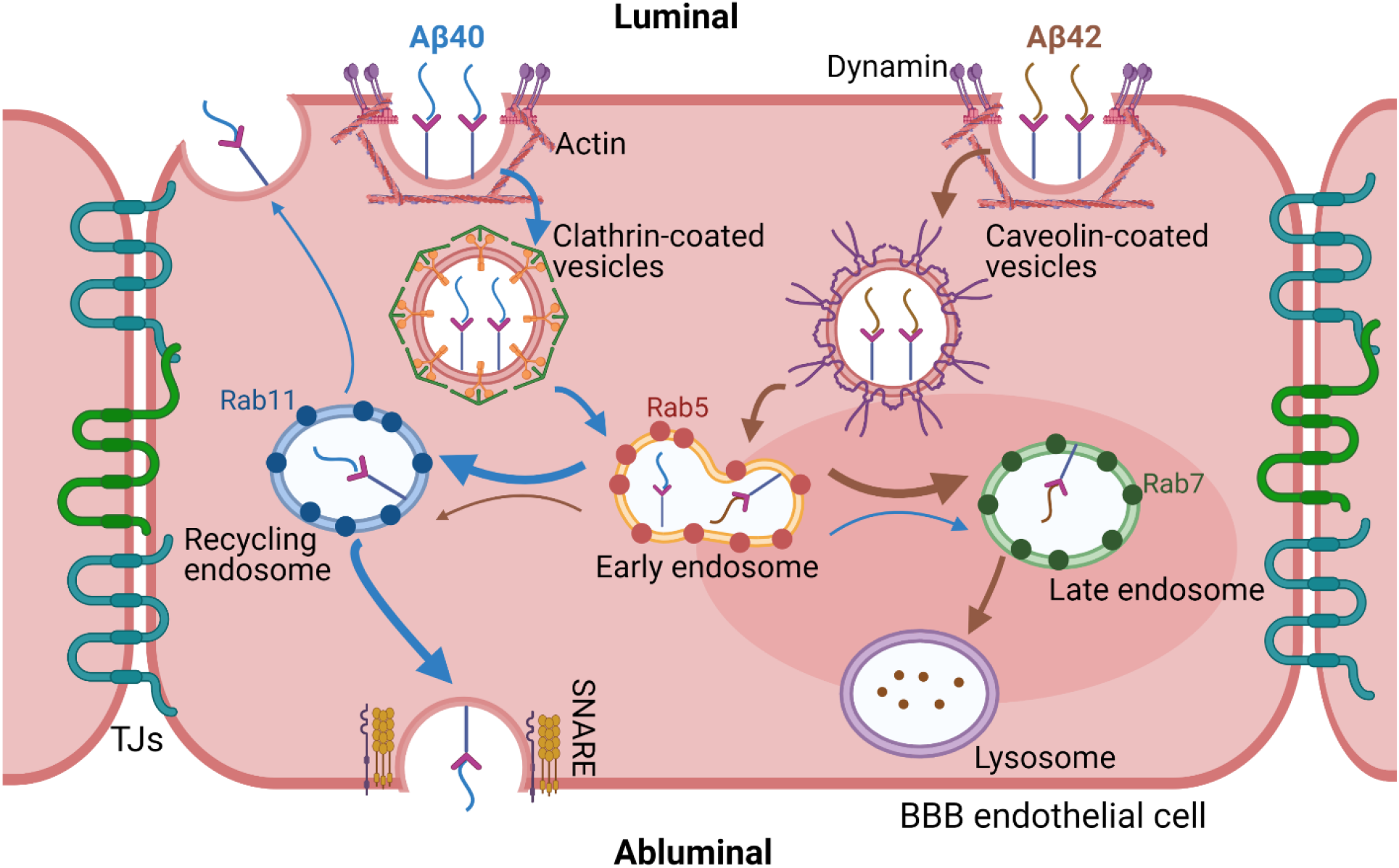
Summary of Aβ40 and Aβ42 endocytosis and intracellular trafficking pathway in the BBB endothelial cells. Both peptides were internalized via energy, dynamin and actin-dependent endocytosis. Endocytosis of Aβ40 was found to be clathrin-mediated whereas Aβ42 was majorly dependent on caveolae. Following endocytosis, the peptides were sorted by the endo-lysosomal system of endothelial cells. Aβ42 was found to accumulate more in the late endosomes and lysosomes (brown thick arrow) but Aβ40 was more likely to get transcytosed to the abluminal side (blue thick arrow) via recycling endosome and soluble N-ethylmaleimide sensitive fusion protein attachment protein receptors (SNARE) protein facilitated exocytosis.

## MATERIALS AND METHODS

### Cell culture

Human brain endothelial (hCMEC/D3) cells were grown according to the culture conditions reported previously^61^. Polarized hCMEC/D3 cell monolayers were cultured on collagen (Corning, MA) coated coverslip bottomed dishes (Mattek, MA), on 6 well plates, or on Transwell® filters (Corning Costar^™^, MA) at 5 % CO_2_ and 37 °C. All the monolayers were moved to low-serum (1 % serum) medium, one night before the experiment. Bovine brain microvascular endothelial (BBME) cells were obtained from Cell Applications Inc. (San Diego, CA). The BBME cellular monolayers were grown as reported previously^62^. Formation of monolayers on Transwell^®^ filters with good tight junctional integrity was confirmed by measuring the trans-endothelial electrical resistance (TEER) across the monolayers.

### Microscopy and cellular imaging

The hCMEC/D3 or BBME monolayers were treated with 1 μM of F-Aβ40, F-Aβ42, Sulforhodamine101 (SR101) labeled Aβ40, or SR101-Aβ42 for 60 min followed by 20 μg/mL of AlexaFluor labeled transferrin (AF633-TRF) treatment for 30 min. In case of inhibitor experiments, the cells were pretreated with inhibitor for the designated time and then incubated with F-Aβ. Then the cells were washed with PBS and the nuclei were stained by incubating with Hoechst dye (0.5 μg/mL in PBS) for 5 min. At the end, the dishes were fixed in 4 % para-formaldehyde (PFA) at 4 °C for 60 min; mounted with ProLong® gold antifade reagent (Life technologies, OR); and dried overnight before imaging by laser confocal microscopy as described in our previous publication^14^.

### Clathrin/caveolin-1 knockdowns and K44-negative dominant dynamin transfections

Clathrin and caveolin-1 knockdown in polarized hCMEC/D3 cell monolayers was performed using siRNA kit containing RNAi mix (Invitrogen, CA) and reduced serum medium, Opti-MEM^™^ (Gibco, NY). The cells were allowed to recover for 48 h in hCMEC/D3 medium (5 % FBS) before the uptake experiment were performed. GFP and non-GFP K44-negative-dominant dynamin transfections of hCMEC/D3 cell monolayers were performed using Lipofectamine reagent (Invitrogen, CA) as per manufacturer’s protocol. The cells were allowed to recover for 24 h in hCMEC/D3 medium (1 % FBS) and the experiment was performed the next day.

### Rab protein transfections

m-Cherry fluorescent protein fused to Rab5 (m-CFP/Rab5), green fluorescent protein fused to Rab7 (GFP/Rab7) and green fluorescent protein fused to Rab11 (GFP/Rab11) plasmids were obtained as described in our previous publications^25^. Transfections of hCMEC/D3 cell monolayers were performed using Lipofectamine reagent (Invitrogen, CA) as per manufacturer’s protocol. The cells were allowed to recover for 24 h in hCMEC/D3 medium (1 % FBS) and the experiment was performed the following day.

### Flow cytometry

Following the treatment with 1 μM F-Aβ for 60 minutes, hCMEC/D3 cells were washed thoroughly with PBS and gently trypsinized with trypsin-EDTA for 2 minutes and neutralized with FBS. In case of inhibitor experiments, the cells were pretreated with inhibitor for the designated time and then incubated with F-Aβ. The dislodged cells were washed twice using ice cold PBS, fixed with 4 % PFA solution and analyzed for intracellular fluorescence using BD FACSCalibur^™^ flow cytometer. The F-Aβ40 and F-Aβ42 intracellular fluorescence intensities were measured using 488 nm laser fitted with 530/30 filter. Data was acquired with BD CellQuest^™^ Pro and analyzed using FlowJo software. The F-Aβ uptake was represented as histograms of intracellular fluorescence intensities. All sample analyses were performed within one hour from the completion of the experiment.

### Western blotting

Following the treatment with 0.25 μM Aβ40 or Aβ42 in DMEM for 15 minutes, the cells were washed three times with PBS and lysed with a RIPA buffer containing protease and phosphatase inhibitors (Sigma-Aldrich, St. Louis. MO). Total protein concentration in the lysates was determined by bicinchoninic acid (BCA) assay (Pierce, Waltham, MA). Lysates (25 μg protein per lane) were loaded onto 4-12% Criterion XT precast gels and proteins were separated by SDS-PAGE under reducing conditions (Bio-Rad Laboratories, Hercules, CA). The proteins were then electroblotted onto a 0.45 μm nitrocellulose membrane. Membranes were blocked with 5% nonfat dry milk protein (Bio-Rad Laboratories, Hercules, CA), followed by overnight incubation at 4 °C with primary antibodies (1:1000) against: phospho-caveolin-1 (Y14) (#3251), caveolin-1 (#3267), Vinculin (#13901) (Cell Signaling Technology, Danvers, MA). Afterwards, the membrane was incubated with IR-dye conjugated secondary antibody (1:2000) for 1 h at room temperature. Immuno-reactive bands were then imaged (Odyssey CLx; LI-COR Inc, Lincoln, NE) and the band intensities were quantified by densitometry (Image StudioTM Lite Software, LI-COR Inc, Lincoln, NE).

### ELISA

Following the 48-hour recovery of vehicle/siRNA transfections, hCMEC/D3 cells were incubated with 1 μM Aβ40/42 for 30 minutes. Afterwards, cells were washed thoroughly with ice-cold PBS twice and harvested in 0.25% trypsin-EDTA. The resulting cell pellets were lysed in 50 μL RIPA buffer containing protease inhibitor. The cell lysates were centrifuged at 10,000 rpm for 10 minutes to remove cell debris and used as samples for ELISA. Aβ levels were determined with Aβ40-and Aβ42-specific ELISA kits (<u>KHB3481</u> and KHB3544, ThermoFisher) according to the manufacturer’s instructions. For data analysis, Aβ concentration was normalized by the total protein concentration and fold change compared to untreated control was calculated. The same cells lysates were also subjected to western blotting as described above in order to confirm the successful knockdown of caveolin-1.

### Live cell imaging

Following the incubation with various fluorophores the cells were washed twice, maintained in an atmosphere humidified with 5% CO2 in air, and imaged live using a TE-2000-S inverted microscope (Chiyoda-ku, Tokyo 100-8331, Japan) equipped with Nikon FITC HQ and m-Cherry-A-zero filters. The images were captured using Nikon’s NIS elements AR 3.0 software and processed using ZEN Imaging software.

### Mechanisms of F-Aβ40 and F-Aβ42 endocytosis at the BBB endothelium

#### 1 Energy dependence of F-Aβ uptake

The polarized hCMEC/D3 cell monolayers grown on Transwell^®^ filters and 6-well culture plates were pre-incubated with glucose free DMEM supplemented with 50 mM 2-deoxy glucose and 0.1 % sodium azide (ATP depletion medium) for 30 min before start of the experiment. The control cells were incubated with regular DMEM medium containing glucose. Then, 1 μM of either F-Aβ40 or F-Aβ42 was added to the luminal side of the Transwell^®^ filter and incubated for 30 min. The resultant intracellular fluorescence was assessed by flow cytometry.

#### 2 Dynamin, actin, clathrin and lipid raft mediated F-Aβ endocytosis

Small molecule inhibitors were used to inhibit the dynamin dependent vesicle pinch-off, actin polymerization, clathrin-dependent endocytosis, or lipid raft-mediated uptake to assess the contribution of these processes to the internalization of FAβ40 and F-Aβ42 by the BBB endothelium. In these experiments, polarized hCMEC/D3 monolayers were pre-incubated with medium containing 1 % serum for at least one hour before the addition of inhibitors. The hCMEC/D3 cells grown on 6-well culture plates or on Transwell® filters were pre-incubated with inhibitors for 30 min, then 1 μM of F-Aβ40 or F-Aβ42 was added and the monolayers were incubated for 60 min. The uptake of F-Aβ peptides was determined using flow cytometry. The following inhibitors and concentrations were used: 80 μM Dynasore (Tocris bioscience, MO) (dynamin inhibitor), 10 μM cytochlasin A (Cayman Chemical, MI) (actin inhibitor), 200 μM monodansylcadaverine (MDC, specific clathrin inhibitor), 10 mM chlorpromazine (MP Biomaterials, OH) (CPZ, non-specific clathrin inhibitor), 2.5 μM nystatin (specific lipid raft inhibitor), 5 mM methyl-β-cyclodextrin (Acros Organics, NJ) (mβCD, non-specific cholesterol chelator). All the control experiments were performed in a similar fashion without the addition of inhibitors and the intracellular fluorescence was assessed by flow cytometry. For knockdown studies, control or knock-down cells were treated with F-Aβ peptides and AlexaFluor 633-transferrin (AF633-Trf) for 60 minutes. The confocal images were obtained after washing the cells with PBS and fixing with 4 % PFA for 1 h at 4 °C.

In a separate experiment, polarized BBME cell monolayers were also pre-incubated with 10 mM mβCD for 30 min, followed by incubation with 1 μM of F-Aβ40 or F-Aβ42, and 20 μg/mL Alexafluor® labeled Transferrin (AF633-TRF) for 60 min. The z-series confocal micrographs were obtained after washing the filters with PBS and fixing with 4 % PFA for 1 h at 4 °C. Then Transwells® were mounted with Vectashield ® Antifade mounting medium with DAPI (Vector Laboratories, CA).

### Intracellular itineraries of F-Aβ40 and F-Aβ42 at the BBB endothelium

### 1 Accumulation in Early Endosomes

hCMEC/D3 cells expressing m-CFP/ Rab5 were incubated with 1 μM F-Aβ40 for 15 minutes. Live cell imaging was conducted up to 60 minutes after washing the cells with DMEM.

### 2 Accumulation in Secondary Endosomes

The cells were treated with 1 μM F-Aβ40 and Dil-LDL (15 μg/ml) for 60 minutes. Then live cell imaging was conducted after washing the cells with DMEM.

### 3 Accumulation in Late Endosomes

hCMEC/D3 cells expressing GFP labeled Rab7 were incubated with 1 μM SR101-Aβ40 or SR101-Aβ42 for 15 minutes. Then the cells were washed with DMEM and imaged by live cell imaging up to 60 minutes.

### 4 Accumulation in Recycling Endosomes

hCMEC/D3 cells expressing GFP/Rab11 were incubated with 1 μM SR101-Aβ40 or SR101-Aβ42 for 15 minutes. Then the cells were washed with DMEM and imaged by live cell imaging up to 60 minutes.

## 5 Accumulation in Lysosomes

The cells grown on coverslip-bottom dishes were treated with 1 μM F-Aβ40 or F-Aβ42 and 75 nM Lysotracker Red® (Invitrogen-Molecular Probes, Carlsbad, CA) for 30 min at 37 °C. Thereafter, the cells were washed with ice-cold PBS 3 times and imaged by confocal microscopy. The fluorescein and Lysotracker Red signals in the image were superimposed, and the extent of colocalization was estimated by Pearson’s correlation coefficients.

## Acknowledgements

This study was supported by the National Institutes of Health National Institute on Aging (AG058081) and National Institute of Neurological Disorders and Stroke (R01NS125437).

